# Npac Is a Co-factor of Histone H3K36me3 and Regulates Transcriptional Elongation in Mouse ES Cells

**DOI:** 10.1101/2020.07.16.205989

**Authors:** Sue Yu, Jia Li, Guanxu Ji, Zhen Long Ng, Jiamin Siew, Wan Ning Lo, Ying Ye, Yuan Yuan Chew, Yun Chau Long, Wensheng Zhang, Ernesto Guccione, Yuin Han Loh, Zhi-Hong Jiang, Henry Yang, Qiang Wu

## Abstract

Chromatin modification contributes to pluripotency maintenance in embryonic stem cells (ESCs). However, the related mechanisms remain obscure. Here, we show that Npac, a “reader” of histone H3 lysine 36 trimethylation (H3K36me3), is required to maintain mouse ESC pluripotency since knockdown of *Npac* causes mouse ESC differentiation. Depletion of Npac in mouse embryonic fibroblasts (MEFs) inhibits reprogramming efficiency. Furthermore, our Npac ChIP-seq results reveal that Npac co-localizes with histone H3K36me3 in gene bodies of actively transcribed genes in mESCs. Interestingly, we find that Npac interacts with p-TEFb, RNA Pol II Ser2 and Ser5. Depletion of Npac disrupts transcriptional elongation of pluripotency genes *Nanog* and *Rif1*. Taken together, we propose that Npac is essential for transcriptional elongation of pluripotency genes by recruiting of p-TEFb and interacting with RNA Pol II Ser2 and Ser5.

## Introduction

Embryonic stem (ES) cells derived from the inner cell mass of the early embryo are characterized by self-renewal and pluripotency, the ability to differentiate into many different cell types [1, 2]. Since ES cells can be cultured indefinitely in vitro, they are a promising resource for regenerative therapy, in particular ES cells show potential for treating degenerative diseases such as diabetes and Parkinson’s disease [3, 4]. Moreover, induced pluripotent stem cells (iPSC) showed enormous potential for the application and progress in gene therapy and regenerative medicine [5-8]. Therefore, enhanced understanding of molecular mechanisms regulating ES cell identity would be of great value toward developing ES and iPSC-based therapies.

Transcription factors Oct4 (encoded by *Pou5f1* gene), Sox2 and Nanog constitute the core transcriptional network that activates genes that promote pluripotency and self-renewal and inhibit genes that promote differentiation [9-12]. Yamanaka’s discovery that the combination of transcription factors OKSM (Oct4, Klf4, Sox2 and c-Myc) was sufficient to reprogram terminally differentiated cells to pluripotent stem cells further proved the importance of those core transcription factors [7]. Aside from these, many other transcription factors are essential for pluripotency [13-16].

Besides transcription factors, chromatin regulators also contribute to mESC pluripotency through providing the necessary environment for proper gene expression [17]. Recently, a handful of chromatin regulators that are critical for ES cell pluripotency were characterized. ES cells contain structurally relaxed and transcriptionally permissive chromatin that allows for epigenetic remodeling [18]. However, factors that modify this epigenetic configuration are not completely known. Lysine-trimethylation modifications at histone H3 are the most stable epigenetic marks on histones. ESCs are featured by higher level of histone H3 lysine 4 trimethylation (H3K4me3), which is generally correlated with gene activation [19]. Conversely, H3K27me3 and H3K9me3 are related to gene silencing and heterochromatin in ESCs [20]. However, few studies have examined the regulation of histone H3K36me3 in ES cells [21, 22]. Histone H3K36me3 marks active genes and preferentially occupies exons and introns (gene bodies) [23] and is considered as a marker of transcriptional elongation. Recently, a large-scale methyl lysine interactome study discovered proteins that bind to specific histone marks [24]. Interestingly, all proteins that bind to histone H3K36me3 have a common PWWP domain. This and other studies [25-27] suggest the essential role of the PWWP domain in binding to histone H3K36me3.

Npac (also known as NP60 and Glyr1) containing a PWWP domain is one of the proteins that can bind to histone H3K36me3 [24]. ChIP-seq analysis on human chromosome 22 revealed that both histone H3K36me3 and Npac are exclusively localized at gene bodies [24], suggesting that Npac may function in transcriptional elongation. Additionally, Npac is a co-factor of LSD2 which mediates histone H3K4 demethylation [28-30]. These findings suggest that Npac regulates gene expression through interacting with specific histone modifications. However, how Npac plays its function is largely unknown.

In this study, we found that Npac is required to maintain mouse ES cell pluripotency. Depletion of Npac leads to mESC differentiation with loss of pluripotency. Depletion of Npac also reduces the reprogramming efficiency of MEFs. We observed that Npac positively regulates pluripotency genes such as *Pou5f1* and *Nanog*. Npac may prevent mESC differentiation by repressing the MAPK/ERK pathway. Furthermore, ChIP-seq experiments showed that Npac co-localizes with histone H3K36me3 in the body of gene which are actively transcribed in mESCs. Npac interacts with RNA Pol II (including Ser2 and Ser5 phosphorylated RNA Pol II) and p-TEFb, and Npac depletion causes transcriptional elongation defect of *Nanog* and *Rif1*. Together, these results establish that Npac maintains mESC pluripotency and regulates transcriptional elongation in mESCs.

## Results

### Npac is required for maintenance of mouse ESC pluripotency

To test whether Npac is associated with ESC pluripotency, we induced mouse ES cells to differentiate by using ES medium without LIF (leukemia inhibitory factor). We observed that *Npac* mRNA level was decreased during differentiation, dropping to around 40% at 5 days after LIF removal (Fig. 1A).

**Figure 1.**
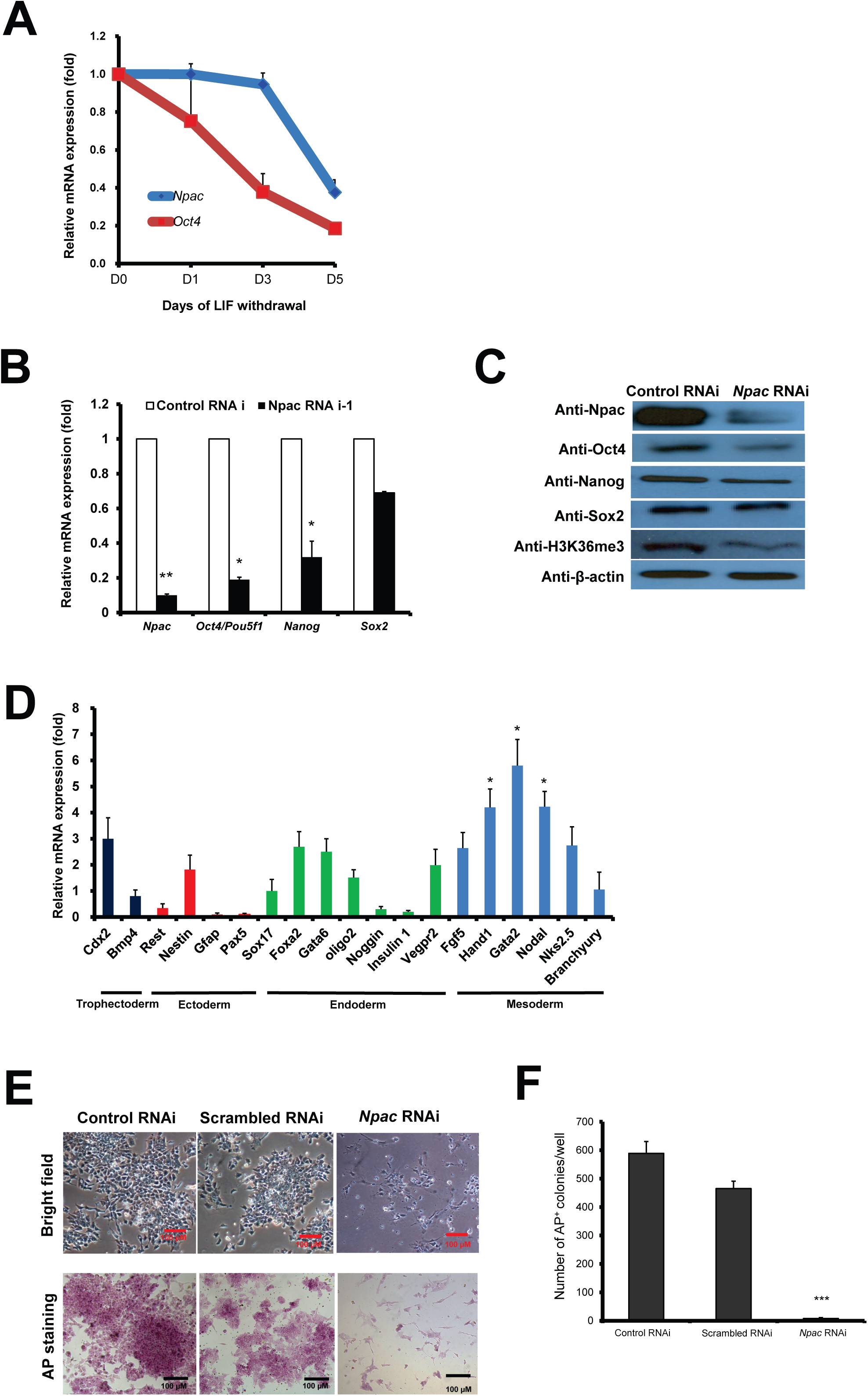
Npac is required to maintain mESC pluripotency. **A**. *Npac* mRNA level was decreased in mES cells cultured in LIF withdrawal ESC medium. The level of the *Npac* and *Oct4* mRNA were compared to control cells cultured in normal ESC medium and normalized against *β-actin*. **B**. Levels of pluripotency genes *Oct4, Sox2* and *Nanog* were significantly decreased upon depletion of Npac. Two different shRNAs targeting distinct regions of *Npac* (*Npac* RNAi-1 was shown in Figure 1B and *Npac* RNAi-2 was shown in Supplementary Figure S1A) were transfected into mESCs to knockdown *Npac*. mESCs transfected with empty pSUPER.puro vector were used as control. **C**. Knockdown of *Npac* resulted in decreased protein levels of Oct4, Sox2, Nanog and histone H3K36me3. β-actin served as loading control. **D**. Depletion of Npac caused up-regulation of lineage specific markers for endoderm and mesoderm. **E**. Representative bright field images (upper panel) of E14 cells transfected with control RNAi, scrambled RNAi or *Npac* RNAi followed by 4 days puromycin selection. Alkaline Phosphatase (AP) staining was shown at the bottom panel. AP staining was conducted on the fourth day of transfection. **F**. Quantification of AP-positive colonies for control RNAi, scrambled RNAi or *Npac* RNAi transfected E14 cells. Scale bar = 100 μm. Specific primers were used to measure gene expression levels by real-time PCR. Gene expression levels were normalized against *β-actin*. All error bars are mean ± SE (n=3). Significance: * P <≤ 0.05, ** P ≤ 0.01, *** P ≤ 0.001.

Next, we depleted Npac by RNAi to determine the role of Npac in ESC pluripotency. Transfection of mouse ES cells with shRNA plasmids (*Npac* RNAi-1 or RNAi-2) targeting *Npac* significantly reduced the level of *Npac* mRNA (Fig. 1B, Supplementary Fig. S1A). We observed that transfection with *Npac* RNAi-1 reduced the mRNA levels of ES cell pluripotency genes *Pou5f1* and *Nanog* significantly (Fig. 1B). Additionally, protein levels of Oct4 and Nanog were also reduced in *Npac* RNAi-1 transfected ES cells as well as histone H3K36me3 (Fig. 1C). *Pou5f1* and *Nanog* mRNA expression levels were similarly down-regulated in *Npac* RNAi-2 transfected ES cells (Supplementary Fig. S1A). The reduction of Oct4 and Nanog upon *Npac* knockdown suggested that Npac depletion may cause loss of ES cell pluripotency since Oct4 and Nanog are master regulators required to maintain ESC pluripotency [13-15,31]. This was further supported by the evidence that expressions of lineage marker genes were up-regulated upon loss of Npac: trophectoderm marker *Cdx2* showed 3 fold increment, endoderm markers *Foxa2, Gata6* and *Vegfr2* displayed 2.7, 2.5 and 2 fold increment respectively, while mesoderm markers, *Nodal, Hand1* and *Gata2* increased by 4.2, 4.2 and 5.8 fold respectively (Fig. 1D). Moverover, *Npac* RNAi transfected ES cells showed morphological differentiation and weaker alkaline phosphatise (AP) activity compared to control RNAi transfected cells, while scrambled *Npac* transfected cells appeared to have similar AP activity as control (Fig. 1E,1F), further indicating that Npac depleted cells were undergoing differentiation.

In order to ensure the *Npac* RNAi was specific, we performed *Npac* RNAi rescue experiment. To construct shNpac-resistant Npac mutant plasmid, specific primers were designed and the plasmid of full-length *Npac* cDNA inserted in pCAG-Neo vector was used as template. We transfected E14 cells with control RNAi or *Npac* RNAi plasmid and selected with puromycin. We then transfect the cells with *Npac* RNAi-resistant plasmid to for RNAi rescue. The cells were selected with neomycin for three days, followed by alkaline phosphatise (AP) staining. We observed susained expression level of pluripotency genes *Pou5f1, Nanog* and *Sox2* and clearly more AP positive cells in rescue treatment (*Npac* RNAi-immune OE) than control, showing that *Npac* RNAi-immune OE cells were resistant to *Npac* RNAi (Supplementary Fig. S1B, S1C). We observed that the changes of ALP staining and pluripotency gene experssion can only be partially rescued. This is like due to *Npac* RNAi resulted in ES cell differantion before we transfected *Npac* RNAi-immune OE plasmid into the cells.

To further confirm the important role of Npac in pluripotency maintenance, we generated emboryoid bodies (EBs) from Npac-depleted cells and control cells. We then cultured the EBs in absence of LIF in low-attachment dishes for 14 days. EBs partially mimic *in vivo* embryonic development [32]. We then performed AP staining and found that both EBs generated from control RNAi cells and *Npac* RNAi cells lost pluripotency, while *Npac* RNAi derivated EBs were much smaller than control group, suggesting that Npac depleted EBs grew more slowly than control EBs (Supplementary Fig. S1D). We also performed qRT-PCR to determine the expression levels of lineage markers in EBs generated from control RNAi and *Npac* RNAi cells (Supplementary Fig. S1E). We found that *Npac* expression level in *Npac*-KD EBs was lower compared to control group. The levels of several mesoderm markers (*Hand1, Gata2* and *Nkx2*.*5*) were much higher in Npac-depleted EBs than that in control EBs. In addition, endoderm markers (*Sox17, Foxa2* and *Vegpr2*) showed higher level in *Npac*-KD EBs compared to control EBs. These results suggest that the depletion of Npac may drive ES cells to differentiate into endoderm and mesoderm lineages, which is consistent with the result of *Npac* knockdown in ES cells shown in Figure 1D.

Having seen the effect of Npac depletion, we next examined whether overexpression of Npac affected ES cell pluripotency and differentiation. To this end, we performed EB formation assay using Npac overexpressing cells. After EB induction, EBs were collected at 7th and 14th day which mimic early and late development respectively. We found that *Npac* was expressed at higher level in the embryoid body at the 7th (Supplementary Fig. S1F) and the 14th day (Supplementary Fig. S1G) in Npac overexpressing EBs. Interestingly, *Oct4* and *Nanog* expression levels were about 2 folds in Npac overexpressing EBs compared to normal EBs, suggesting that pluripotency genes were sustained longer in Npac overexpressing EBs. Also, we found the size of Npac OE EBs was bigger than that of control group, suggesting that Npac overexpression may promote EBs to grow faster than control group (Supplementary Fig. S1H).

Based on these results, we conclude that Npac is required to maintain ESC pluripotency. On one hand, Npac depletion represses pluripotency genes and activate lineage marker genes. On the other hand, pluripotency gene expression is maintained upon Npac overexpression in differentiating ES cells.

### Reprogramming efficiency of MEFs to iPSCs is reduced upon Npac depletion

Because of the essential role of Npac in mouse ES cell pluripotency, we next tested its role in reprogramming of somatic cells. *Pou5f1*-GFP MEFs were used to facilitate identification of putative iPSC colonies based on GFP expression [33]. We first confirmed that *Npac* relative expression was decreased to about 29% of the control with OKSM only when MEFs were infected with OKSM along with *Npac* knockdown virus (Fig. 2A). We found that GFP^+^ colonies produced by OKSM plus *Npac* knockdown was 3.5 fold less than the control 14 days later after infection (Fig.2B). We also confirmed this by checking iPSC colonies with AP staining (Fig. 2C, 2D). In addition, we performed immunostaining to examine whether the iPSCs generated from OKSM plus *Npac* knockdown induction were pluripotent. We found that those iPSCs expressed endogenous Oct4 and Nanog, indicating that they were ES-cell like (Fig. 2E). Further, we generated embryoid bodies from GFP^+^ iPSCs which induced by OKSM+*Npac* KD. Our immunostaining results showed that these iPSCs could express lineage markers of endoderm (Nestin), mesoderm (SMA) and ectoderm (Gata4) (Fig. 2F). These results showed that iPSCs generated from OKSM plus *Npac* knockdown are pluripotent.

**Figure 2.**
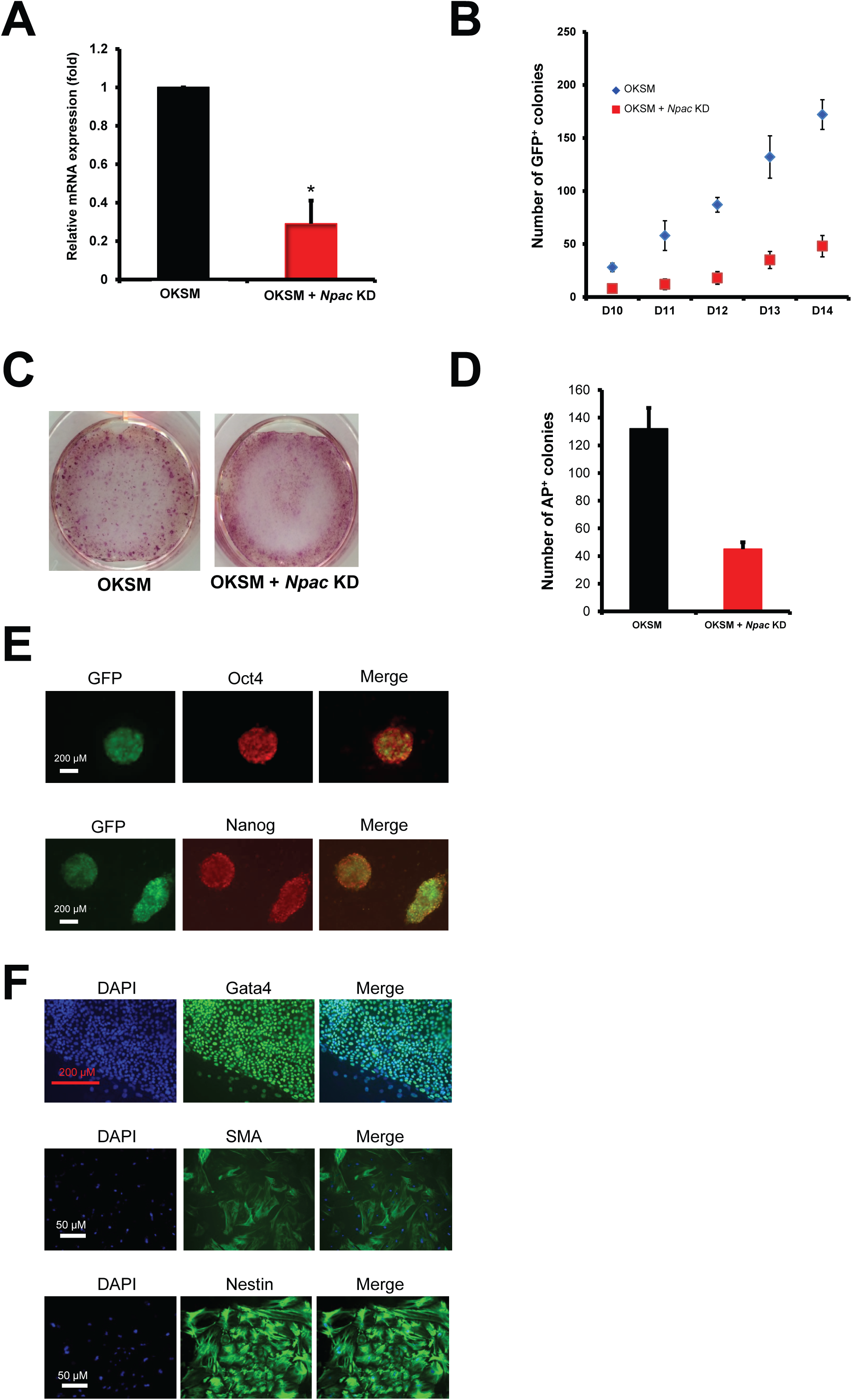
Depletion of Npac inhibits the efficiency of reprogramming. **A**. Npac mRNA relative expression level was determined by real-time PCR in mouse embryonic fibroblasts (MEFs) which were infected with OKSM factors (Oct4, Klf4, Sox2, c-Myc) or OKSM + Npac knockdown retrovirus. RNA was extracted from cells which were harvested 4 days after virus infection. Gene expression levels were normalized against *β-actin*. **B**. Depletion of Npac inhibited reprogramming efficiency process. The numbers of GFP^+^ colonies which indicate putative iPSCs were counted from day 10 to day 14 after virus infection. GFP^+^ colonies formed by OKSM factors + Npac knockdown virus were lower than OKSM control throughout the whole reprogramming process. **C**. The iPSCs generated from OSKM + Npac knockdown virus presented weaker alkaline phosphatase activity than OKSM virus. There were less AP stained colonies generated from OKSM+Npac knockdown compared to OKSM. **D**. Graphical representation of AP staining results was shown in Figure 2**C**. **E**. The iPSCs generated from OKSM plus Npac KD expressed Oct4 and Nanog, indicating that they were ES-cell like. Immunostaining was performed with anti-Oct4 and anti-Nanog antibodies in GFP^+^ iPSCs generated from OKSM+Npac KD. **F**. Embryoid bodies generated from GFP^+^ iPSCs which were induced by OKSM+Npac KD were able to express ectoderm, mesoderm and endoderm lineage markers. Embryoid bodies were stained with anti-Gata4, anti-alpha smooth muscle actin (SMA) and anti-Nestin antibodies. DAPI (blue) served as nucleus marker. Scale bar = 100 μm.

Thus, Npac is essential for not only pluripotency maintenance in mES cells but also generation of iPSCs.

### Depletion of Npac represses pluripotency genes while activating development related genes

We next investigated how Npac functions in pluripotency maintenance by profiling gene expression following shRNA-induced *Npac* knockdown. Global expression altered genes after *Npac* knockdown were shown in Supplementary Table S2. Upon Npac depletion, 2696 genes were increased by >1.5 fold and 891 genes were down-regulated (decreased by >1.5 fold) (Fig. 3A). We randomly chose 10 up-regulated and 9 down-regulated genes and tested by qPCR to confirm the gene expression microarray results (Supplementary Fig. S2A, S2B).

**Figure 3.**
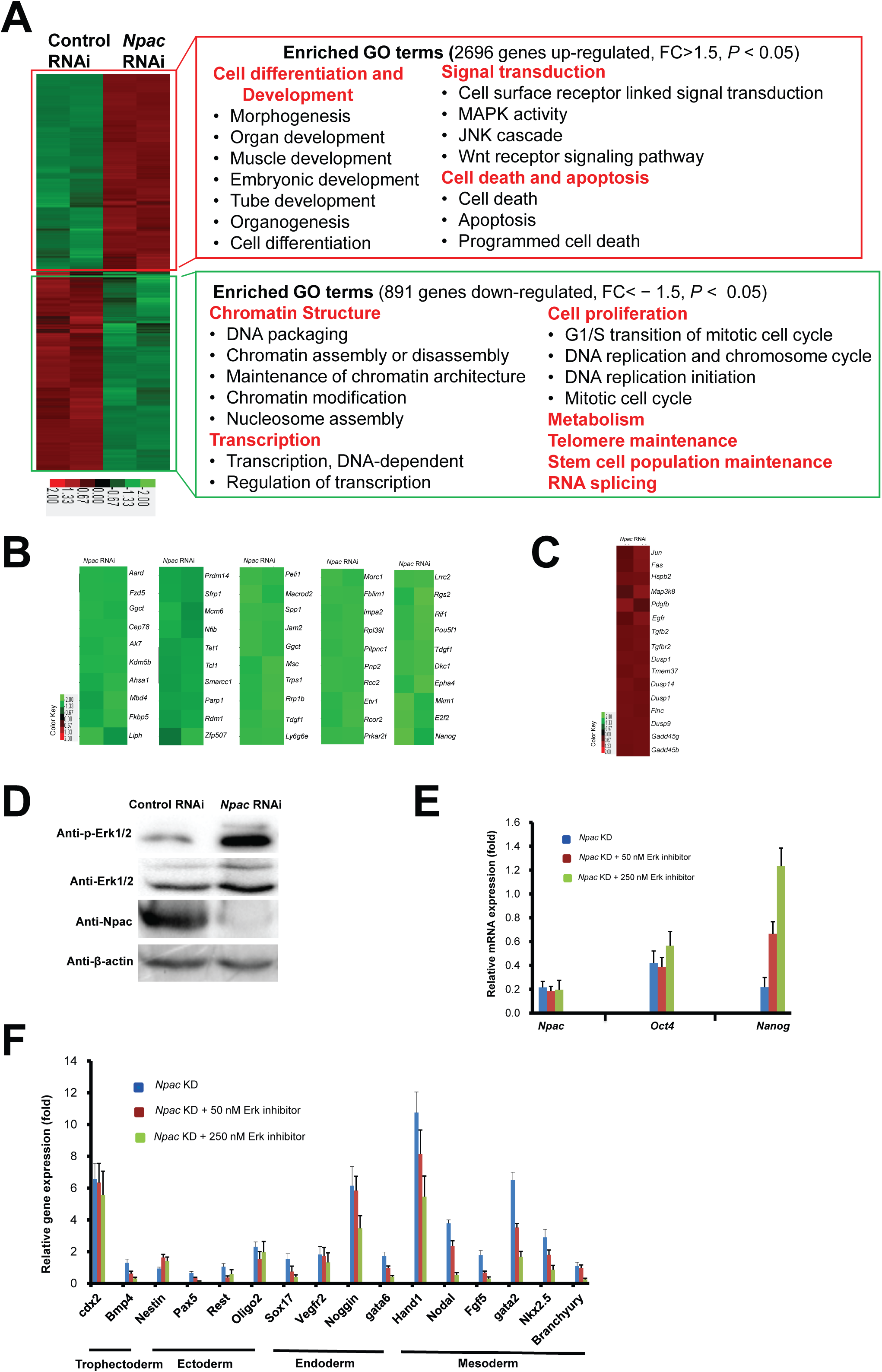
Changes of global gene expression upon Npac depletion in mouse ESCs. **A**. Microarray heat map generated from relative gene expression levels. Relative highly expressed genes were shown in red and low expressed genes in green. *Npac* was knocked down in E14 cells and selected for 96 hours. Then whole genome cDNA microarray hybridization was performed. Duplicates were chosen to ensure reproducibility of results. Gene ontology (GO) analysis was performed relating to “biological process” for the up- or down-regulated genes respectively. The enriched categories were classified into several function groups and listed in the figure. **B**. Heatmap of down-regulated pluripotency genes upon *Npac* knockdown in mESCs. Genes were selected according to their known functions in pluripotency. Each selected gene was taken as individual tiles from the thumbnail-dendogram duplicates. **C**. Heatmap of up-regulated MAPK pathway-related genes upon *Npac* knockdown in mESCs. Genes were selected according to their known functions in MAPK pathway. **D**. p-ERK1/2 and ERK1/2 protein levels were elevated in Npac depleted cells as compared to control cells. β-actin served as loading control. **E**. ERK inhibitor triggered elevated expression of Nanog while it could not rescue the down-regulated expression of master pluripotency gene *Oct4* upon Npac depletion. **F**. ERK inhibitors slightly brought down the up-regulated lineage markers in *Npac* knocked-down cells. Mouse ESCs were transfected with *Npac* RNAi plasmid or control plasmid and 50 nM or 250 nM of ERK inhibitors (PD0325901, Sigma) were added into selection medium for 4 days followed by RNA extraction.

We carried out Gene Ontology (GO) analysis for activated and repressed genes (Fig. 3A). Full list of the enriched terms is shown in Supplementary Table S3. Among genes down-regulated by *Npac* knockdown, enriched categories were related to chromosome modification, suggesting that Npac is required to maintain the unique chromatin structure in mESCs. Notably, we found a majority of known pluripotency genes were down-regulated upon Npac depletion (Fig.3B). Npac could also play important roles in cell proliferation and telomere maintenance, since GO terms related to these were significantly enriched. Among up-regulated genes, many enriched terms were related to development.

### Npac regulates the MAPK/ERK pathway to influence mESC pluripotency

Interestingly, many up-regulated genes upon Npac depletion were linked to Wnt and MAPK signaling pathways that are involved in mESC pluripotency (Fig. 3A, 3C). Nichols et al. reported that suppression of the MAPK/ERK pathway can contribute to the maintenance of mES cell ground state and pluripotency [34, 35]. Also, the ERK pathway promotes mES cell differentiation [36]. We found that levels of ERK1/2 and phosphorylated ERK1/2 (p-ERK1/2) were elevated upon Npac depletion (Fig. 3D). Thus, inhibition of the MAPK/ERK pathway by Npac could contribute to the effect of Npac on pluripotency and differentiation.

To explore the role of the MAPK pathway in Npac function, we tested whether inhibition of the MAPK pathway by ERK inhibitor (PD0325901, Sigma) could rescue the effect of *Npac* knockdown. We first confirmed that 50 nM/250 nM ERK inhibitor (PD0325901, Sigma) was able to block the MAPK pathway in E14 ES cells (Supplementary Fig. S3). We found that the ERK inhibitor did not affect *Npac* knockdown efficiency (Fig. 3E). However, addition of the ERK inhibitor elevated the level of *Nanog*. The ERK inhibitor did not rescue the down-regulation of *Pou5f1* by Npac depletion (Fig. 3E). This is in line with the previous finding that ERK pathway inhibition up-regulates Nanog in ES cells [37, 38]. Also, the addition of the ERK inhibitor reduced the expression levels of lineage markers (Fig. 3F). Finally, Npac depleted cells with or without the ERK inhibitor displayed similar differentiated morphology.

Taken together, our results suggest that Npac depletion activates the MAPK/ERK pathway, leading to mESC differentiation. However, since blocking the MAPK/ERK pathway did not rescue the differentiation phenotype and the down-regulation of *Pou5f1*, Npac likely affects pluripotency also by other unknown mechanisms.

### Npac depletion promotes apoptosis

We also observed that many genes related to cell death and apoptosis were up-regulated when *Npac* was knocked down (Fig. 4A). To evaluate the effect of Npac depletion on cell death, we performed propidium iodide staining and flow cytometry. FACS analysis found that the percentages of cells in sub-G1 phase were significantly increased in Npac depleted cells compared to the control (Fig. 4B, 4C), suggesting that there was a sub-G1 phase arrest in the cell cycle. Furthermore, Annexin V staining assay showed that apoptotic cells increased to 40.3% of the total cell population upon Npac depletion, compared to only 9.9% of the total cell population in the control (Fig. 4D, 4E). These results indicate that depletion of Npac causes apoptosis.

**Figure 4.**
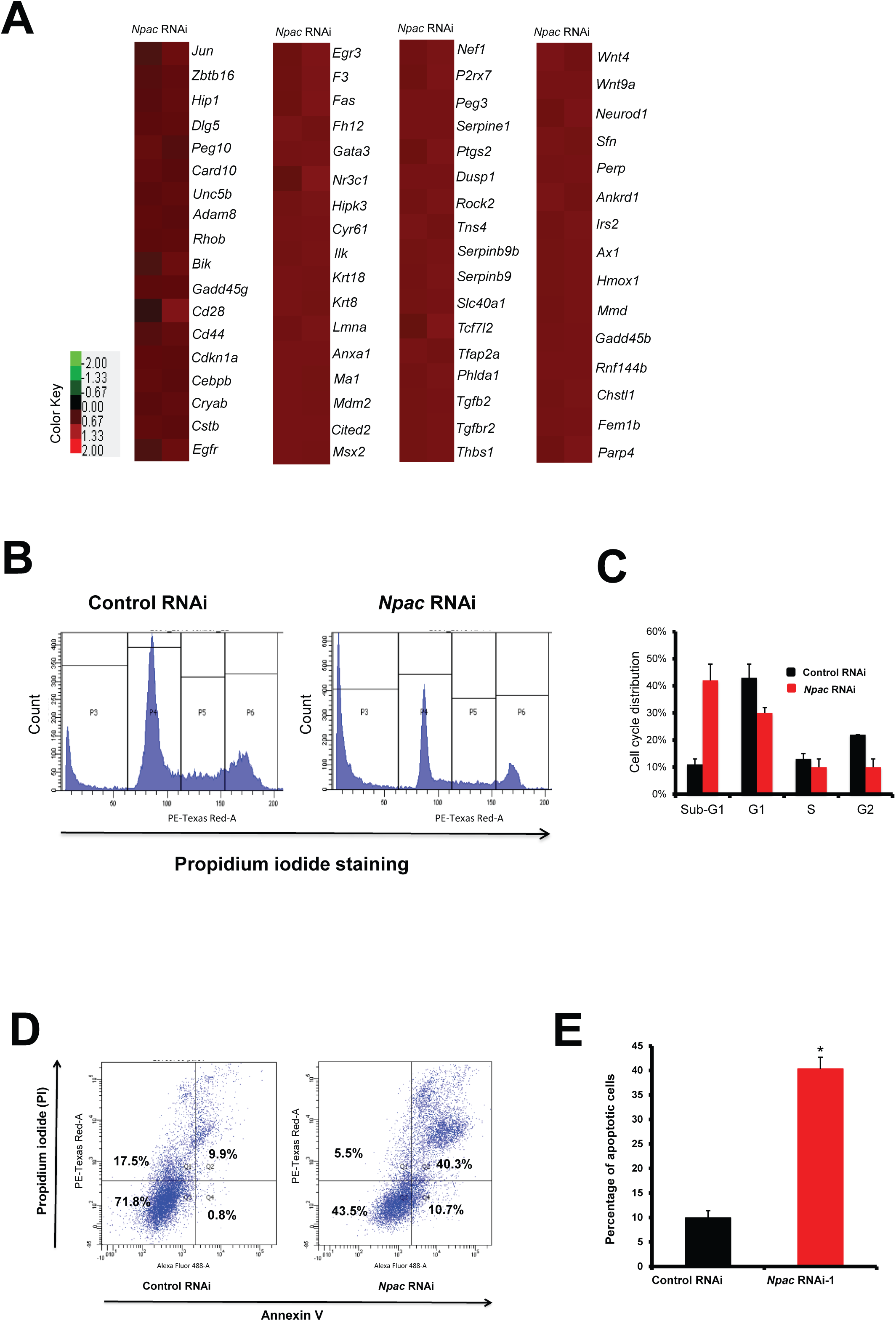
Npac depletion may cause cellular apoptosis. **A**. Heatmap of up-regulated cell death related genes upon *Npac* knockdown in mouse ESCs. Genes were selected according to their known functions in cell death. **B**. Cell cycle analysis by flow cytometry in *Npac* RNAi cells and control RNAi group. **C**. The representative flow cytometry pattern is shown. **D**. Apoptosis triggered by Npac depletion was analyzed by Annexin V staining assay through flow cytometry. **E**. Graphical representation of the apoptosis cells by Annexin V staining assay. Mouse E14 cells were transfected with *Npac* RNAi or empty plasmid as control. After 96 hours selection, cells were harvested for cell cycle analysis or Annexin V staining assay followed by flow cytometry. Error bars were based on three separate experiments.

### Npac is located at gene bodies and co-occupies genomic sites of histone H3K36me3

Oct4 and Nanog are master regulators in the pluripotency transcriptional network [13]. Since depletion of *Npac* down-regulates *Nanog* and *Oct4*, we tested whether Npac is located to *Pou5f1* and *Nanog* promoters using chromatin immunoprecipitation (ChIP). Interestingly, we found enrichment for Npac in introns and exons (here defined as the gene body) of *Nanog* but not the promoter (Fig. 5A, 5B). Also, we did not observe any enrichment within the promoter of *Pou5f1* (Supplementary Fig.S4). Similarly, we also found enrichment in the gene bodies of other pluripotent genes such as *Tcf15, Prdm14* and *Tcl1* (Fig. 5C).

**Figure 5.**
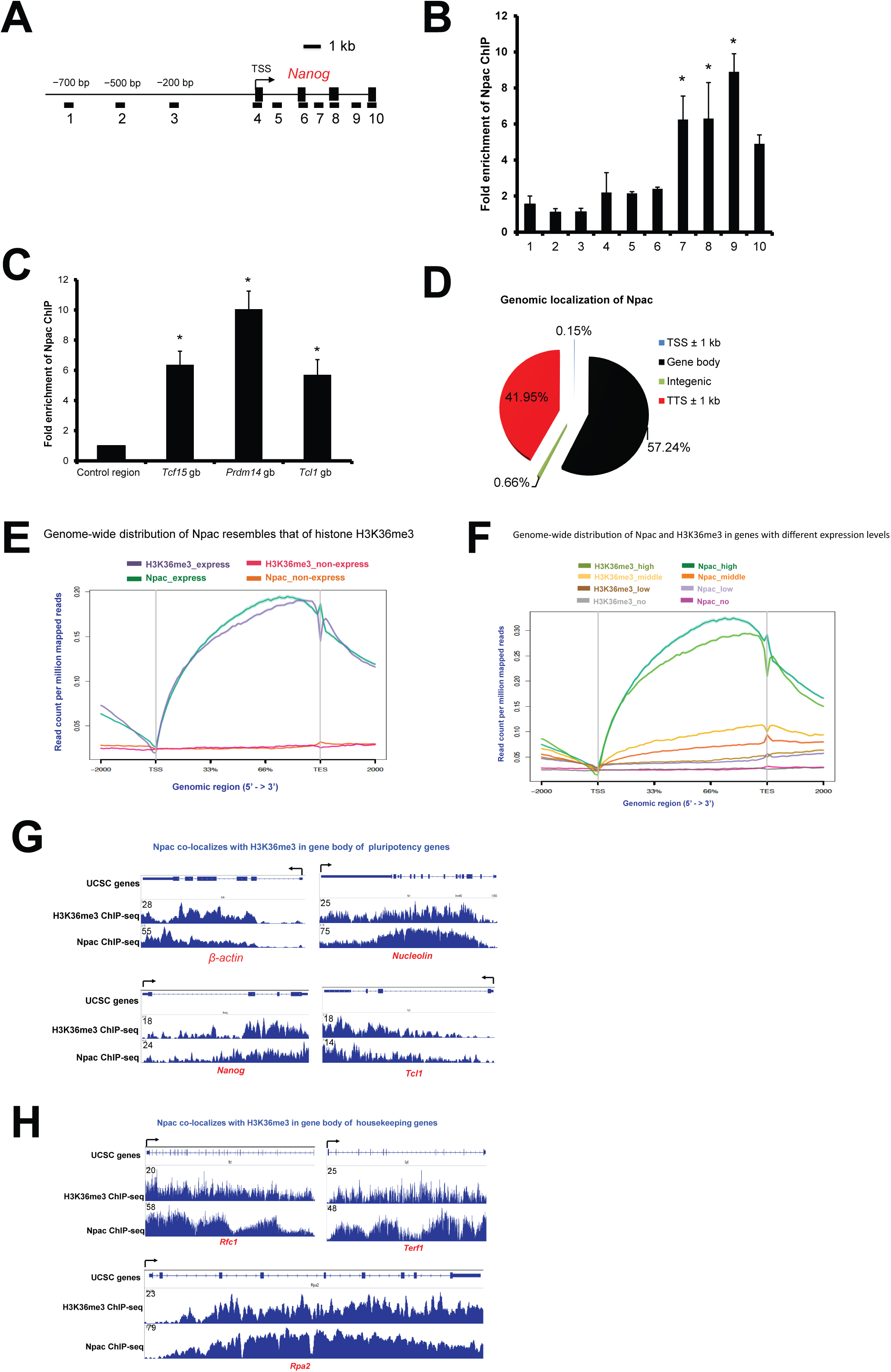
Npac is mainly located to gene bodies and its genome-wide distribution resembles that of histone H3K36me3. **A**. Schematic diagram at the bottom shows the primers designed at specific areas upstream and downstream of *Nanog* gene. **B**. Npac is associated with *Nanog* gene body, with high enrichment fold at gene body of Nanog. Real time PCR primers were designed according to *Nanog* genomic region. **C**. Npac is also associated with gene bodies of other pluripotency genes including *Tcf15, Prdm14* and *Tcl1*. Real time PCR primers were designed at gene bodies of *Tcf15, Prdm14* and *Tcl1* genomic regions. (D) Genome-wide distributions of Npac in mESCs. **E**. Genome-wide distribution of Npac resembled that of histone H3K36me3. Both are enriched at actively transcribed genes while having low or no enrichment in non-expressed genes in E14 cells. H3K36me3_express represents H3K36me3 genome wide distribution in expressed genes in E14 cells; Npac non-express represents Npac genome distribution in non-expressed genes in E14 cells. **F**. Genome-wide distribution of Npac and H3K36me3 in genes with different expression levels (high, middle, low, no). H3K36me3_high (middle, low, no) represents H3K36me3 genome wide distribution in mESC genes with high (middle, low, no) expression. Npac high (middle, low, no) represents Npac genome wide distribution in mESC genes with high (middle, low, no) expression. Each gene body is represented from 0% (transcriptional start site; TSS) to 100% (transcriptional termination site; TTS). **G**. Housekeeping genes (*ACTB*) and pluripotency genes (*Nucleolin, Nanog* and *Tcl1*) representatives of Npac and H3K36me3 ChIP-seq peaks in mESCs. Arrows denote TSS and transcription orientation. **H**. Telomere maintenance related genes (*Rfc1, Terf1* and *Rpa2*) representatives of Npac and H3K36me3 ChIP-seq peaks in mESCs. Arrows denote TSS and transcription orientation.

To determine the genome-wide distribution of Npac in ES cells, we conducted a ChIP-seq experiment using anti-Npac antibody. We identified 12414 potential genomic sites of Npac where 2416 genes were mapped, of which 57.24% sites were located within gene bodies (the global binding sites of Npac were shown in Supplementary Table S4). Additionally, 41.95% of the sites were within transcription termination sites (TTS), followed by 0.66% and 0.15% respectively mapped to intergenic and transcription start sites (TSS) (Fig. 5D). Gene ontology analysis showed the genes that Npac binds to are linked to development, transcription, chromatin modification, cell cycle and RNA processing (Supplementary Table S5).

Since Npac is a cofactor of LSD2 which demethylates histone H3K4me1 and H3K4me2, we were also interested in the relationship between Npac and histone H3K4me2. Indeed, we found that the genome-wide profile of Npac localization was inversely correlated to that of histone H3K4me2 (Supplementary Fig. S5A). We were also keen to explore whether Npac is linked to other histone modifications. Here, we chose several important histone modifications (histone H3K9me3, histone H3K27me3 and histone H3K4me3) (Supplementary Fig. S5B) and their respective modifiers (Eset, Ezh2 and MLL2) (Supplementary Fig. S5C), as well as ESC-enriched transcription factors (TFs) (Oct4, Nanog and Sox2) (Supplementary Fig. S5D) to compare their binding with that of Npac. We found that Npac binding profile displayed unique pattern compared to those epigenetic modifiers, histone modification and ESC-enriched TFs. In general, Npac-associate genes (most are active genes) are much less than that of others. Further, Npac, unlike Eset, Ezh2 and MLL2 (mainly located at TSS sites), is enriched in gene bodies and 3’ ends. Thirdly, Npac shares some genomic loci with master TFs Oct/Nanog/Sox2 but the genomic locations of these three TFs are clearly different from that of Npac.

Npac is a putative reader of histone H3K36me3 together with which Npac are present almost exclusively over gene bodies [24]. We found that in mESCs, the genome-wide distribution of Npac resembled that of histone H3K36me3, and both of them were enriched at expressed genes in E14 ES cells, which displayed absence from the transcription start sites (TSS), but gradually increased from gene bodies to transcription termination sites (TTS), while had low or even no binding at inactive genes in E14 ES cells (Fig. 5E). Further, we separated genes into 4 groups (high, middle, low and no) according to their gene expression levels. The results also showed similar genome-wide distribution between Npac and H3K36me3, which displayed most enriched at genes with high expression, but lower binding in genes with lower expression (Fig. 5F). This further indicates that Npac and H3K36me3 are enriched in actively transcribed genes in E14 ES cells. Indeed, we observed that both histone H3K36me3 and Npac had high occupancies in actively transcribed genes, such as housekeeping gene *ActinB*, pluripotency genes *Nanog, Nucleolin* and *Tcl1* (Fig. 5G), and telomere maintenance related genes *Rfc1, Terf1* and *Rpa2* (Fig. 5H). On the other hand, we observed clearly low Npac and histone H3K36me3 occupancies on inactive genes. These genes included developmental genes (Supplementary Fig. S6A), MAPK pathway related genes (Supplementary Fig. S6B) and cell death related genes (Supplementary Fig. S6C). Taken together, these results show that Npac co-localizes with histone H3K36me3 in gene bodies of active genes in mES cells, suggesting that Npac plays roles in histone H3K36me3-associated cellular functions including gene activation and transcriptional elongation.

### Npac is likely involved in transcriptional elongation

Next, we examined how Npac is involved in transcriptional elongation. We found that Npac can interact with RNA Pol II (Fig. 6A). This result is in line with the previous report that LSD2 complex may include Pol II and Npac [28]. Next, we found that Npac can also associate with Ser2 and Ser5 phosphorylated RNA pol II (Fig. 6B, 6C). In addition, we found that phosphorylation levels of Ser2 and Ser5 were down-regulated upon Npac depletion, while RNA Pol II expression was not affected (Fig. 6D). This suggests that the interaction of Npac with phosphorylation Ser2 and Ser5 may affect their expression and function. Given that Ser5 phosphorylation is associated with transcriptional initiation and early elongation while Ser2 phosphorylation correlates with transcriptional elongation [39], we propose that Npac affects transcriptional elongation through associating with RNA Pol II phosphorylation Ser2 and Ser5.

**Figure 6.**
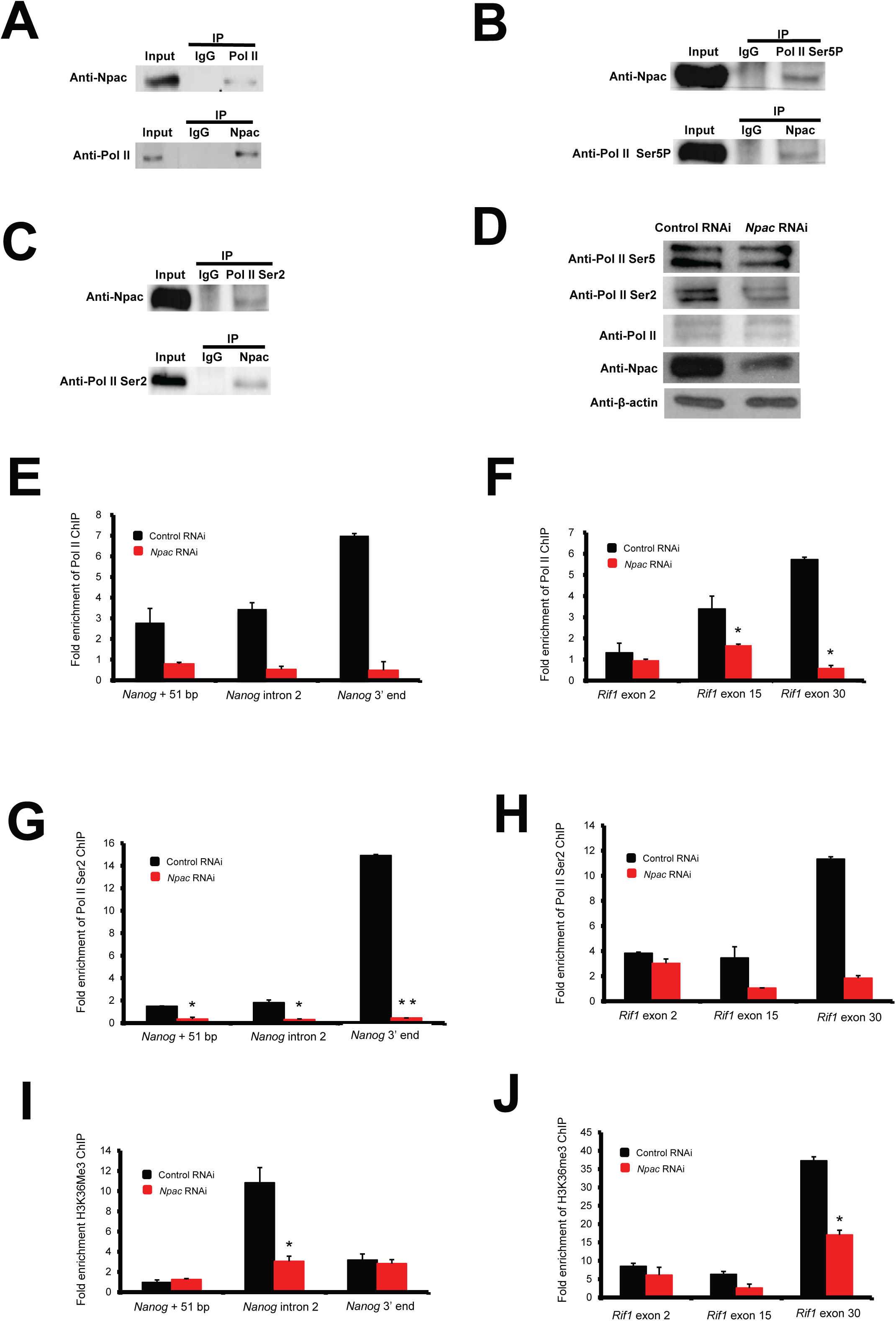
Npac could be involved in RNA Pol II transcriptional elongation. **A**. Npac interacted with RNA Pol II. Cell lysate of wild type ESCs was immunoprecipitated using either anti-Npac antibody or anti-RNA Pol II antibody. Western blot was subsequently carried out with anti-RNA Pol II antibody or anti-Npac antibody. **B**. Npac can be pulled down with RNA Pol II Ser5P. **C**. Npac was associated with RNA Pol II Ser2. **D**. Npac depletion led to down-regulation of RNA Pol II Ser2 while protein levels of RNA Pol II Ser5 and total RNA Pol II were not affected. β-actin served as loading control. The binding of RNA Pol II at gene bodies of *Nanog* (**E)** and *Rif1* (**F)** were significantly reduced in *Npac* depleted cells. The binding of RNA Pol II Ser2 at gene bodies of *Nanog* (**G)** and *Rif1* (**H)** were significantly reduced in *Npac* depleted cells. Histone H3K36me3 binding to pluripotency genes *Nanog* (**I)** and *Rif1* (**J)** was significantly reduced compared to control.

In order to determine whether Npac is essential for RNA Pol II elongation in mouse ESCs, we performed ChIP with RNA Pol II, RNA Pol II Ser5 and RNA Pol II Ser2 in Npac depleted cells and control cells. We performed ChIP-qPCR for the gene bodies of two pluripotency genes *Nanog* and *Rif1*, and *Utrn*, a gene up-regulated in Npac depleted cells. We found that the presence of RNA Pol II and RNA Pol II Ser2 at gene bodies of *Nanog* (Fig. 6E, 6G) and *Rif1* (Fig. 6F, 6H) was significantly reduced in *Npac* knockdown cells, while their presence at *Utrn* (Supplementary Fig. S7A, S7B) was not significantly changed. In addition, the level of H3K36me3 was also reduced at gene bodies of *Nanog* (Fig. 6I) and *Rif1* (Fig. 6J) in Npac depleted cells. However, the binding of RNA Pol II Ser5 at gene bodies of *Nanog* (Supplementary Fig. S7C) and *Rif1* (Supplementary Fig. S7D) was similar between Npac depleted cells and control cells, suggesting RNA Pol II Ser5 binding is independent of Npac. Taken together, these results suggest that Npac promotes transcriptional elongation. But it does not affect transcriptional initiation.

### Npac associates with p-TEFb to promote transcriptional elongation

Next, we observed that Npac could interact with transcriptional elongation factor b (p-TEFb), which is composed of Cyclin T1 and Cdk9 (Fig. 7A, 7B). p-TEFb can phosphorylate the carboxyl-terminal domain (CTD) of the large subunit of RNA polymerase II, thus promoting transcriptional initiation and elongation [40]. Thus, Npac could act as an essential component of the elongation complex. To test whether Npac is required for transcriptional elongation, we performed elongation recovery assay to measure the recovery of transcription at different positions of two genes (*Nanog* and *Rif1*). We incubated ESCs with 100 uM 5,6-Dichloro-1-β-D-ribofuranosylbenzimidazole (DRB) which is widely used as an elongation inhibitor [41]. After three hours, ESCs were washed twice with PBS and cultured with fresh medium before total RNA was isolated in every 5 minutes (Fig. 7C) [37]. We first confirmed that *Npac* knockdown efficiency was not affected by addition of DRB in Npac depleted cells, which showed about 40% *Npac* mRNA compared to the control (Fig. 7D). Next, we examined transcripts from *Nanog* and *Rif1* at different positions after release from the elongation block. Following the release, transcriptional output at the exon 1 of *Nanog* and exon 2 of *Rif1* was not significantly affected by *Npac* RNAi (Fig. 7G, 7I). However, in Npac-depleted cells, the recovery of transcription at downstream regions (exon 4 region of *Nanog* and exon 30 of *Rif1*) was significantly reduced compared to the control (Fig. 7F, 7J). Taken together, these results suggest that Npac depletion causes transcriptional elongation defect of *Nanog* and *Rif1*.

**Figure 7.**
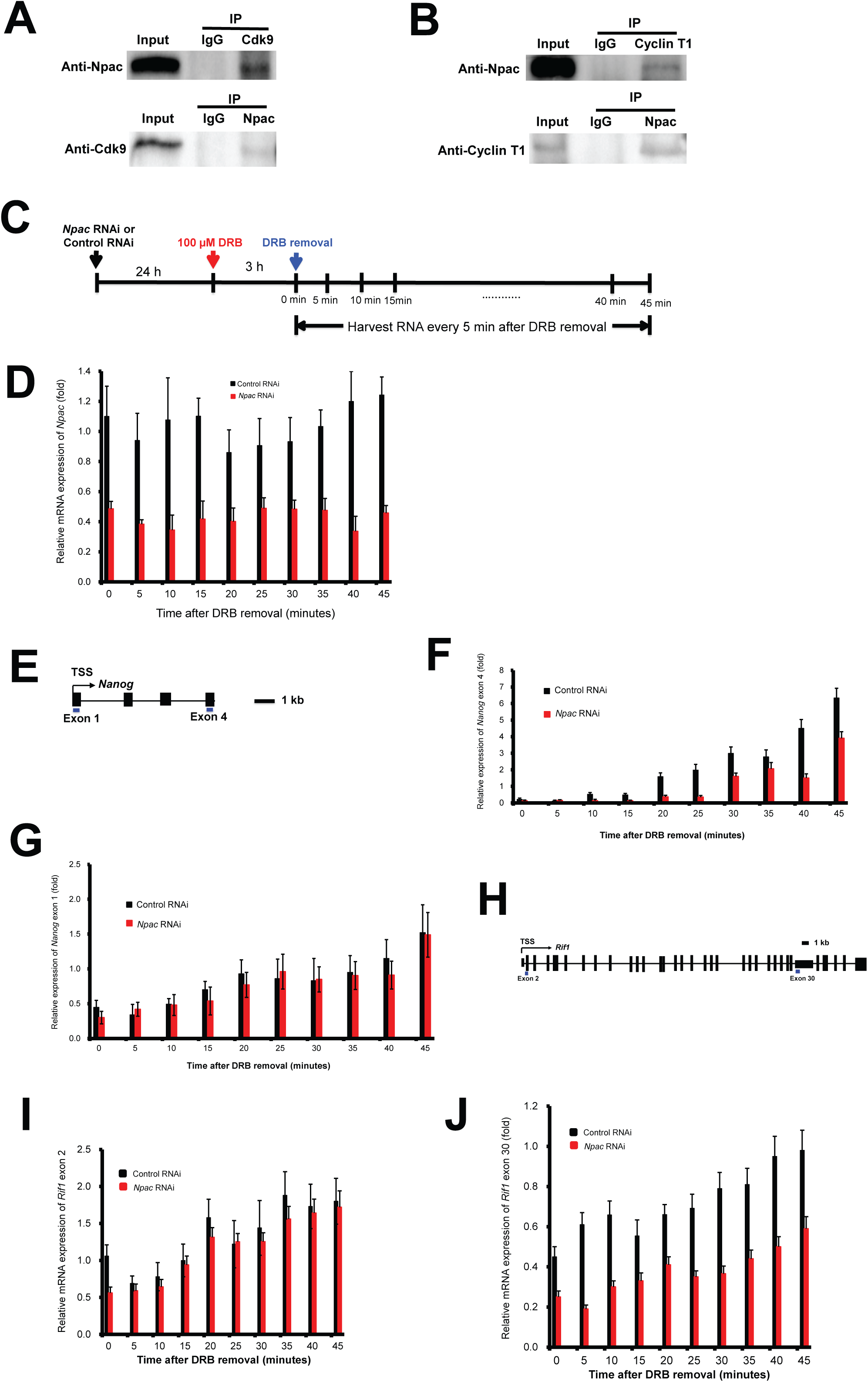
Npac interacts with positive transcriptional elongation factor (p-TEFb) and Npac depletion may lead to transcriptional elongation defect. **A**. Npac interacts with Cdk9. **B**. Npac is associated with Cyclin T1. Control IP was performed using anti-IgG antibody. **C**. Elongation recovery assay process. After E14 cells were transfected with *Npac* RNAi or control RNAi for 24 hours, 100 μM DRB was applied and followed by 3 hours incubation, washed twice with PBS and incubated in fresh medium before total RNA was extracted in every 5 minutes. **D**. *Npac* knockdown efficiency was not affected with the addition of DRB in Npac depleted cells. Npac mRNA expression was detected by real time PCR and expression levels were normalized against *β-actin*. **E**. The analyzed regions of *Nanog* were shown schematically in the map in E. (**F)** & (**G**) Changes in the rate of transcription of different portions of *Nanog* upon depletion of Npac. **H**. The analyzed regions of *Rif1* were shown schematically in the map in (H). (**I**) & (**J**) Changes in the rate of transcription of different regions of *Rif1* upon depletion of Npac. The recovery of transcription was assessed at different positions. Each graph illustrates the RNA levels at different regions of *Rif1* at different recovery time after DRB block was released. The graphs represent averages of three independent experiments and standard deviations were provided.

## Discussion

Mouse ESC pluripotency is governed by both genetic and epigenetic mechanisms. Many pluripotency factors including transcription factors and epigenetic regulators have been discovered in ES cells [42]. Our results indicate that Npac is required to maintain pluripotency in mouse ES cells. First, we found that depletion of Npac significantly repressed expression of master pluripotency factors Oct4 and Nanog. Besides these core factors, many other known ESC pluripotency factors were also decreased upon Npac depletion according to gene expression microarray. Among these, *Tet1* is specifically expressed in ESCs and required for ES cell maintenance [43]. *Tcl1*, a cofactor of the Akt1 kinase, is essential for self-renewal of ES cells [44]. Also, KDM5B, a histone H3 trimethyl lysine 4 (H3K4me3) demethylase, is an activator of ESC self-renewal correlated genes [45]. Second, transient knockdown of Npac increased expression of mesoderm and endoderm linage markers and reduced alkaline phosphatase activity. These results further support the assertion that Npac is required for maintaining ES cell in an undifferentiated state. Third, we found that loss of Npac activated the MAPK signaling pathway (Fig. 3A, 3D). ERK signalling pathway can induce ES cell differentiation into all germ layers *in vitro* [46, 47]. In addition, activation of ERK represses Nanog expression and causes ES cell differentiation into primitive endoderm [48]. It is intriguing that *Npac* knockdown leads mESCs to differentiation but ERK inhibitors did not fully rescue the differentiation phenotype. It is noteworthy that ERK inhibitors can block general ESC differentiation and thus may mask true differentiation defects of Npac-depleted ESCs. Thus ERK inhibitors might not be specific to rescue the phenotype resulted from Npac depletion. Therefore, though it is possible that reduction of *Nanog* upon Npac depletion was partially caused by the activation of ERK pathway, this is unlikely to be the sole mechanism. We surmise that Npac depletion also results in changes in chromatin state, RNA-binding and cell metabolism, some of which may be non-reversible. It is highly likely that Npac regulates pluripotency using some other unknown mechanisms which will be interesting to be further explored.

Furthermore, the function of Npac in somatic reprogramming verified its essential role in pluripotency. There are several possible ways in which Npac depletion inhibits reprogramming process. Reprogramming consists of a set of molecular processes that transform a somatic cell into a pluripotent stem cell. In the process of reprogramming, genes related to differentiated state should be repressed first and markers associated with pluripotency will be activated subsequently. Meanwhile, widespread chromatin remodelling occurs during the whole process [49, 50]. Our microarray results showed that Npac depletion activates many development associated genes (Fig. 3A) and down-regulates a subset of pluripotency genes. Thus, reprogramming may be blocked initially by high expression of somatic genes. It has been demonstrated that active marks such as H3K4me2/3 can cause chromatin to be in “open” state and thus enhance iPSC formation, while repressive marks such as H3K9me and H3K27me function in opposite way and impair iPSC formation [50]. Given that H3K36me3 is a mark classically associated with active transcription, we can predict that active H3K36me3 marks may promote reprogramming. Thus the fact that *Npac* knockdown causes down-regulation of H3K36me3 (Fig. 1C) could be a reason why iPSC formation is inhibited. In addition, GO analysis of microarray and ChIP-seq revealed that Npac targets many genes that are associated with chromatin modification and nucleosome assembly (Fig. 3A). Therefore, this provides additional evidence that Npac depletion may impair reprogramming by inhibiting permissive chromatin state. Finally, inhibition of the ERK pathway not only enables maintenance of mouse ES cells in ground pluripotent state, but also can enhance somatic reprogramming [35, 51]. Therefore, lower reprogramming efficiency in *Npac* knockdown MEFs could also be affected by activation of the ERK pathway.

We found that Npac depletion can increase cellular apoptosis (Fig. 4B, 4D). The mechanisms of apoptosis have been elucidated in numerous studies, classically includes both extrinsic and intrinsic pathways [52]. Among them, the ERK pathway and p53-dependent apoptosis are two important mechanisms. ERK activity can boost apoptotic pathways by activation of caspase-8 [53]. Thus, the fact that Npac depletion increases expression of ERK (Fig. 3D) and *caspase-8* (gene expression microarray results) could explain apoptosis caused by loss of Npac. p53 causes apoptosis by transcription-dependent and independent mechanisms [54]. Therefore, apoptosis could be induced by affecting p53 downstream targets. Indeed, we observed an up-regulation of *PERP* expression in microarray (Fig. 4A), which is a proapoptotic gene targeted by p53 [55]. APAF-1 is another protein up-regulated upon Npac knockdown according to our microarray result. APAF-1 is a component of the apoptotic machinery activated by p53. Because of the unique abbreviated cell cycle of mouse ES cells, mESCs display a different mechanism of cell cycle arrest and apoptosis compared to somatic cells. Our results showed that Npac-depleted ES cells arrest in the sub-G1 phase (Fig. 4B), this could be another reason that ES cells undergo apoptosis upon Npac depletion. Taken together, our results suggest that Npac depletion causes apoptosis through the p53-dependent pathway and the ERK pathway.

We found that Npac is co-localized with transcriptional elongation mark histone H3K36me3 in gene bodies of actively transcribed genes in ESCs (Fig. 5E, 5F). This is consistent with the finding from a study in human Hela cells [24]. However, it remains unclear whether recruitment of Npac depends on the localization of histone H3K36me3. Given that Npac predominantly occupies actively transcribed genes (Fig. 5G, 5H), it appears that Npac functions as a transcriptional activator of those actively transcribed genes (such as pluripotency genes and telomere maintenance genes) in mouse ES cells. Moreover, we observed that global level of RNA Pol II Ser2 was reduced, while total RNA Pol II was unaffected when Npac was depleted (Fig. 6D). There is a possibility that the lower level of phosphorylated RNA Pol II in Npac depleted cells makes elongation process slower or even blocks transcriptional elongation, therefore most of the active genes in mouse ES cells were down-regulated upon Npac depletion. In addition, the binding of RNA Pol II and RNA Pol II Ser2 at pluripotency genes *Nanog* and *Rif1* were significantly reduced upon Npac depletion, while the binding of RNA Pol II Ser5 at these two genes was not significantly changed. These further confirmed that Npac is required for transcriptional elongation.

In mammalian cells, Ser2 of RNA Pol II can be phosphorylated by the Cdk9 kinase subunit of p-TEFb, which results in transiting elongation initiation to productive elongation [56]. According to previous studies, some specific activators, such as DNA or RNA bound activators and co-activators can recruit p-TEFb to transcription units.

For example, one chromatin remodelling protein Brd4 recruits p-TEFb to stimulate RNA polymerase II-dependent transcription [57]. Given the association of Npac with p-TEFb in our study (Fig. 7A, 7B), it is possible that Npac may recruit p-TEFb to chromatin and further results in successful elongation of transcription of target genes. This is in concert with the observation that Npac depletion caused transcriptional elongation defect for pluripotency genes *Nanog* and *Rif1*. However, this may not be the sole reason. Lower enrichment of histone H3K36me3 and RNA Pol II Ser2 upon Npac depletion may also contribute to transcriptional elongation defect. Taken together, these imply an essential role of Npac in elongation process.

Interestingly, according to our Npac gene expression microarray results, we found there are more upregulated than down regulated genes upon Npac depletion. This appears to be contrary to the fact that Npac is associated with actively transcribed genes. However, we think *Npac* knockdown in ESCs results in differentiation and triggers significant upregulation of abundant developmental genes which are silenced in undifferentiated ESCs. During the differentiation process, active pluripotency genes become inactivated meanwhile silenced developmental genes are activated. Since the number of activated developmental genes is bigger than that of inactivated pluripotency genes, there are more upregulated genes than downregulated genes upon Npac depletion. Further, the activation of MAPK pathway caused by *Npac* knockdown will trigger many differentiation-related genes. Last but not least, the upregulation of developmental genes in *Npac* knockdown cells is probably independent of Npac-mediated transcription elongation and it could be triggered by ES cell differentiation. Nevertheless, given that Npac may have diverse functions, it is of interest to further explore how Npac plays its role in gene regulation during ES cell differentiation.

In summary, we propose a model that Npac regulates mESC pluripotency and influence transcriptional elongation by interaction with p-TEFb, RNA Pol II Ser2 and Ser5 (Fig.8A, 8B).

**Figure 8.**
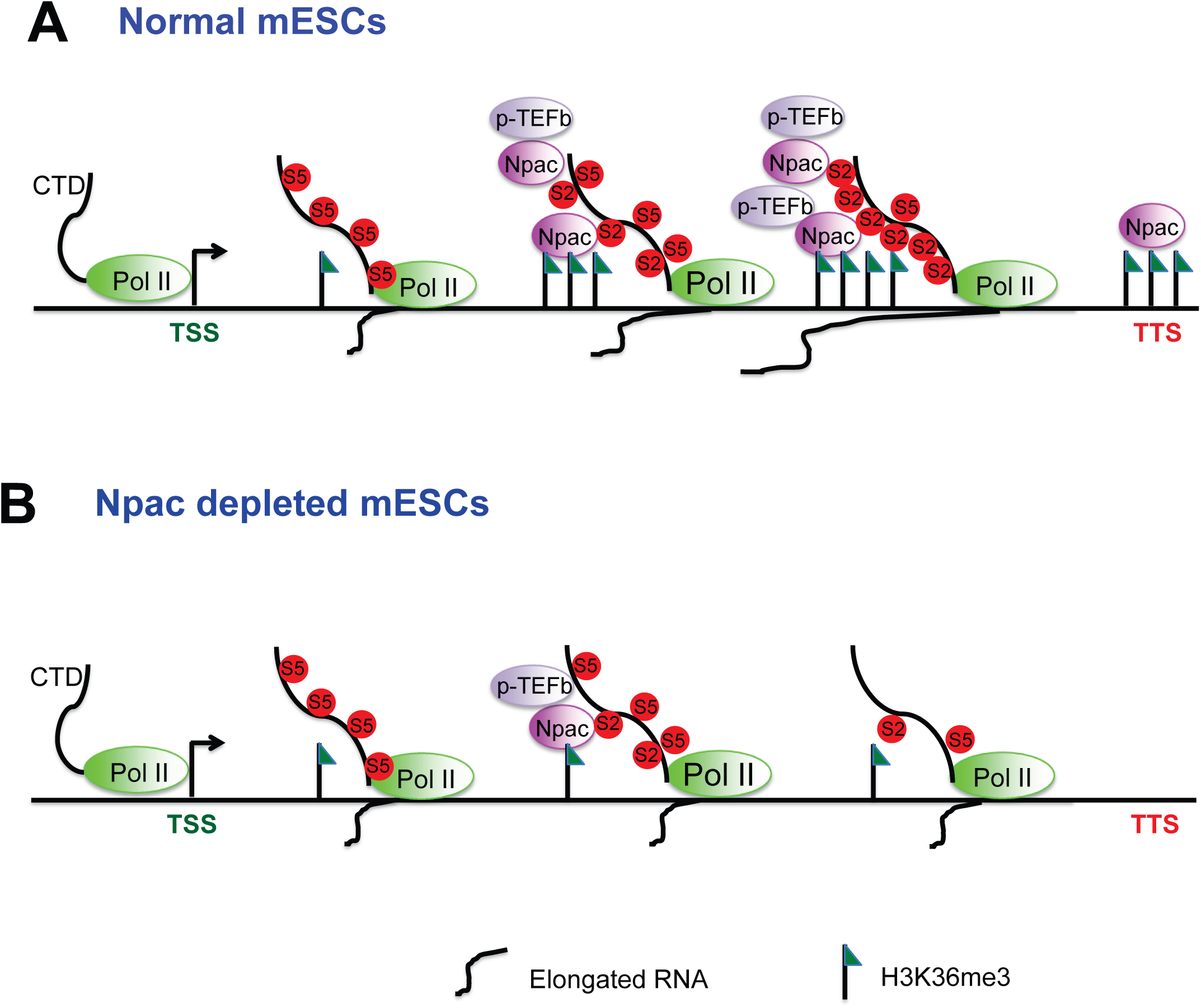
Model depicting the role of Npac in pluripotency. **A**. In normal mouse ES cells, Npac is expressed at high level and histone H3K36me3 is enriched in the gene bodies of actively transcribed genes. Npac interacts with RNA Pol II Ser2 and recruits p-TEFb to promote productive elongation. **B**. In Npac depleted cells, reduction of Npac leads to reduced enrichment of Pol II Ser2 and histone H3K36me3 at pluripotency genes, thus blocking productive transcriptional elongation.

## Material and methods

### Cell culture

In this study, mouse E14 ES cells (ATCC® CRL-1821™, Manassas, VA), SNL feeder cells (CBA-316, Cell Biolabs. San Diego, CA), Platinum-E cells (Plat-E) (RV-101, Cell Biolabs. San Diego, CA) and *Oct4*-*GFP* mouse embryonic fibroblasts (MEFs) were cultured in a 37°C CO2 incubator with 5% CO2 as previously described [58].

### Construction of plasmids

Plasmids for shRNA-1 and shRNA-2 targeting *Npac* were designed using Eurofins MWG Operon siMAX™ software. Oligonucleotides were inserted into pSuper.puro vector (VEC-pBS-0008, Oligoengine. Seattle, WA). The primers for Npac overexpression were designed by Primer 5 software to amplify full length cDNA of mouse *Npac*/*Glyr1* (NM_028720.2) and the PCR product was inserted into Bgl II and Mlu I site of pPyCAGIP vector. To construct retrovirus packaging plasmids, full-length *Npac*/*Glyr1* cDNA was ligated into MluI and NotI restriction sites of pMXs plasmid (18656, Addgene. Watertown, MA). To construct Npac mutant plasmid that produces functional Npac protein but was resistant to Npac RNAi targeting, specific primers were designed with silent mutations in protein coding domain sequence. The Npac mutant plasmid was generated according to the manual of Q5^®^ Site-Directed Mutagenesis Kit (E0554S, New England Biolabs. Ipswich, MA). The plasmid of full-length *Npac* cDNA inserted in pCAG-Neo vector was used as PCR template. The sequences of the primers were shown in Supplementary Table 1.

### Transfection, RNA extraction, reverse transcription and real-time PCR

Transfection was conducted using Lipofectamine 2000 (11668019, Invitrogen. Waltham, MA) according to the protocol. Cells were selected by 1 µg/ml puromycin for 4 days after transfection. Either protein or RNA was then extracted from the cells. RNA extraction, reverse transcription and real-time PCR were performed as previously described [58]. Sequences of qPCR primers were shown in Supplementary Table S1.

### Gene expression microarray analysis

Mouse ES cells (E14) were transfected with *Npac* knockdown plasmid or control plasmid and cultured for 4 days with selection. RNA was then extracted from the cells. Gene expression microarray data were analyzed as previously described [58,59]. The microarray results have been deposited to NCBI GEO (GSE93296).

### Chromatin immunoprecipitation (ChIP) assay and ChIP-sequencing (ChIP-seq)

Chromatin immunoprecipitation (ChIP) assay and ChIP-sequencing (ChIP-seq) were conducted as previously described [58,59]. Antibodies used for ChIP were: anti-Npac (14833-1-AP, Proteintech Group. Rosemont, IL), anti-histone histone H3K36me3 (ab9050, Abcam, Cambridge, UK), anti-RNA polymerase II CTD repeat YSPTSPS (phospho S2) (ab5095, Abcam. Cambridge, UK), anti-RNA polymerase II CTD repeat YSPTSPS (phospho S5) (ab140509, Abcam. Cambridge, UK). The ChIP-seq results have been deposited to NCBI GEO (GSE95671).

### Bioinformatics Analysis

Npac ChIP-seq reads were mapped to the mouse genome (NCBI37/mm9) using Burrows-Wheeler Aligner mapping software [60]. After removing duplicate reads, the mapped results identified board peaks with MACS2. For location classification, ChIP-seq peaks were annotated by comparing the locations of all transcription start sites and terminal sites in mouse genome with Perl scripts. (10 kb-1 kb upstream of the TSS site defined as upstream, 1 kb upstream of the TSS to the TSS site defined as TSS, regions between the TSS site and the TTS site defined as gene body, 1 kb downstream of the TTS to the TTS site defined as TTS, 10 kb-1 kb downstream of the TTS site defined as downstream).

H3K36me3 ChIP-seq (ENCSR000CGR), H3K4me3 ChIP-seq (GSM1258237) and its modifier MLL2 ChIP-seq (GSM1258241), H3K9me3 ChIP-seq and its modifier Eset ChIP-seq (GSM440256), H3K27me3 ChIP-seq (GSM1199184 & GSM1199185) and its modifier EZH2 ChIP-seq (GSM1199182 & GSM1199183), Oct4 ChIP-seq (GSE65093), Nanog ChIP-seq (GSM915363) and Sox2 ChIP-seq (GSM1179561) on mouse E14 ES cells downloaded from ENCODE were chosen as datasets to compare with Npac ChIP-seq.

E14 RNA-seq (GSM1276712) was also downloaded from ENCODE. We used STAR software [61] to carry out the RNA-seq mapping with the mm9 genome. By analyzing the mapped RNA-seq data, featureCounts [62] was used to obtain the gene expression of E14 sample. All genes were further separated into two groups based on whether the genes are expressed or not. Genes were also classed into 4 groups based on their expression levels (high, middle, low and no). Expression levels were classed according to the number of reads that mapping to mm9 genome. Reads=0 represents non_express, while reads > 0 represents express. 0< reads ≤ 10 represents low expression, 10< reads ≤ 100 represents middle expression, reads > 100 represents high expression. With the respective gene lists and mapped ChIP-seq files, heatmap and average reads distribution were generated with ngsplot [63].

### Western blot

Western blot was performed as described [58,59]. Primary antibodies used in this study were: anti-Npac (14833-1-AP, Proteintech Group. Rosemont, IL), anti-NP60 (sc-390601, Santa Cruz. Dallas, TX), anti-β-actin (sc-81178, Santa Cruz. Dallas, TX), anti-Oct4 (sc-8628, Santa Cruz. Dallas, TX), and anti-Nanog (sc-33760, Santa Cruz. Dallas, TX), anti-Sox2 (sc-99000, Santa Cruz. Dallas, TX), anti-p-ERK (4370, Cell Signaling. Danvers, MA), anti-ERK (137F5, Cell Signaling. Danvers, MA), anti-histone H3K36me3 (ab9050, Abcam. Cambridge, UK), anti-Pol II (sc-899, Santa Cruz. Dallas, TX), anti-RNA polymerase II CTD repeat YSPTSPS (phospho S2) (ab5095, Abcam. Cambridge, UK), anti-RNA polymerase II CTD repeat YSPTSPS (phospho S5) (ab140509, Abcam. Cambridge, UK), anti-Cyclin T1 (sc-10750, Santa Cruz. Dallas, TX), anti-Cdk9 (sc-484, Santa Cruz. Dallas, TX), anti-mouse IgG (sc-2025, Santa Cruz. Dallas, TX), anti-goat IgG (sc-2028, Santa Cruz. Dallas, TX) and anti-rabbit IgG (sc-2027, Santa Cruz. Dallas, TX).

### Alkaline phosphatase (AP) staining

Alkaline phosphatase (AP) staining was conducted with Alkaline Phosphatase Detection Kit (SCR004, Millipore. Burlington, MA) as described in the manufacturer’s protocol. Axio Observer A1 inverted light microscope (Zeiss. Gottingen, Germany) was used to take pictures for AP staining results.

### Co-immunoprecipitation

Cells were lysed in cell lysis buffer (50 mM Tris-HCl pH 8.0, 1 mM ethylenediaminetetraacetic acid (EDTA), 150 mM sodium chloride (NaCl), 1% NP-40, 10% glycerol) with protease inhibitor (4693159001, Roche. Basel, Switzerland) and rotated for 1 hour at 4°C. After precleared by protein G beads (15920010, Invitrogen. Waltham, MA) for 2 hours at 4°C, the cell lysate was incubated overnight with beads bound by specific antibodies at 4°C. Then beads were washed four times with cell lysis buffer and heated in 2X loading dye for 10 minutes at 95°C. The supernatant was used for Western blotting with specific antibodies. IgG antibody (12-371, Chemicon. Temecula, CA) was used for control IP.

### Retrovirus packaging and infection

Retrovirus packaging and infection were carried out as before [58]. Briefly, pMXs retroviral plasmids or pSUPER.retro.puro plasmids were transfected into Plat-E cells. The cells were selected with 1 μg/ml puromycin (P8833, Sigma-Aldrich. St. Louis, MO) and 10 μg/ml blasticidin (A1113902, Life Technologies. Carlsbad, CA) for 36 to 48 hours. Retroviruses were harvested and concentrated with centrifugal filter units (C7715, Millipore Burlington, MA). *Pou5f1*-*GFP* MEFs were seeded into 24-well plates for 6 hours and then infected with retroviruses. Infected MEFs were seeded onto SNL feeder layers 2 days after infection and cultured with mESC medium without LIF until the 5^th^ day post infection. The MEFS were then cultured with KSR medium from 6^th^ day after infection. Numbers of GFP^+^ colonies were recorded daily until day 14. AP staining assays were also conducted at day 14.

### Annexin V-FITC apoptosis assay

Annexin V-FITC apoptosis assay was carried out as described in manufacturer’s protocol (APOAF, Sigma. St. Louis, MO). After transfected with *Npac* RNAi or control RNAi plasmid in 6-well dishes and selected for 4 days. The cells were stained with Annexin V FITC and propidium Iodide. The cells were then analyzed by flow cytometer (BD FACSCanto. BD Biosciences. San Jose, CA).

### Cell cycle analysis

Cell cycle analysis was conducted as before [58]. Briefly, ES cell were transfected with *Npac* RNAi plasmid or control plasmid and selected with puromycin for 4 days. Then cells were stained with 50 μg/ml propidium iodide and then analyzed by the flow cytometer (BD FACSCanto. San Jose, CA) using Flowing Software 2.5.0.

### Transcription elongation assay

Transcriptional elongation assay was carried out as previously described [64, 65]. E14 cells were transfected with *Npac* RNAi-1 or control RNAi. After 24 hours cells were treated with 100 μM 5,6-dichloro-1-bold beta-D-ribofuranosylbenzimidazole (DRB) (287891, Sigma St. Louis, MO) for 3 hours, washed twice with PBS and cultured in fresh medium for different durations (5 min to 45 min). Total RNA was extracted and qRT-PCR was performed to quantify relative expression level changes at different regions along the *Rif1* and *Nanog* genes. Gene expression levels were normalized against β-actin. Sequences of used primers were listed in Supplementary Table S1.

### Statistical analyses

All experiments were conducted in triplicates. Student’s t-test was applied for statistical analysis and the results were mean±SE. P<0.05 was considered significant. Significance: * P ≤ 0.05, ** P ≤ 0.01, *** P ≤ 0.001.

### Data availability

The *Npac* RNAi microarray results have been deposited to NCBI GEO (GSE93296). The Npac ChIP-seq results have been deposited to NCBI GEO (GSE95671).

## Authors’ contributions

QW and HY conceived and designed the experiments. SY, JL, GJ performed the experiments and analysed the data. ZLN, JS, WNL, YY, YYC, YCL, WZ, EG, YHL and ZHJ contributed reagents/materials/analysis tools. SY, JL and QW wrote the manuscript. SY, WZ and QW revised the paper. All authors read and approved the final manuscript.

## Acknowledgments

We thank Dr. Takao Inoue for critical reading of the manuscript. We thank Professor Huck Hui Ng for providing *Oct4-GFP* MEFs. This work was supported by Singapore National Medical Research Council (CBRG14nov065) and Macau Science and Technology Development Fund (FDCT-18-033-SKL-016A).

## Supplementary material

**Supplementary Figure S1 Npac is essential for mESC pluripotency. A**. mRNA levels of pluripotency genes Oct4, Sox2 and Nanog were slightly repressed upon depletion of Npac with *Npac* RNAi-2. Npac RNAi-2 was transfected into mESCs to knockdown Npac. ESCs transfected with empty pSUPER.puro vector were used as a control. **B**. Representative bright field images (upper panel) and AP staining pictures (lower panel) of *Npac* RNAi rescue experiment. E14 cells were transfected with control RNAi or *Npac* RNAi first and followed by puromycin selection for 2 days. Then those transfected cells were rescued by transfection of Npac-Immune OE plasmid followed by neomycin and puromycin selection for 3 days. Cells transfected with control empty vector were control group. Alkaline phosphatise (ALP) staining was conducted. Scale bar = 100um. **C**. RNA isolation and qRT-PCR were performed to compare gene expression levels of pluripotency marker genes and Npac after *Npac* RNAi rescue. Gene expression levels were normalized against *β-actin*. **D**. Representative bright fields and Alkaline Phosphatase (AP) staining images of EBs generated from E14 transfected with control RNAi or *Npac* RNAi. **E**. Gene expression of lineage markers in EBs was determined by qRT-PCR. Embryoid bodies were generated by culturing them in low-attachment culture plates for 14 days. AP staining was conducted at day 14 and EBs were harvested in 14th day for RNA isolation and qRT-PCR. **F&G**. Pluripotency genes were sustained in Npac OE EBs. **H**. Representative bright fields and AP staining results of EBs generated from control or Npac OE E14 cells.

**Supplementary Figure S2 Validation of *Npac* RNAi gene expression microarray data**. Specific primers were designed to check the respective gene expression levels of randomly selected down-regulated (**A**) and up-regulated (**B**) genes upon *Npac* knockdown in mouse ESCs. Gene expression were normalised against *β-actin*.

**Supplementary Figure S3 Erk inhibitor (PD0325901) is able to block MAPK pathway**. mESCs (E14 cells) were incubated with 50 nM or 250 nM Erk inhibitor for 24 hours and DMSO was added into E14 cells as control. Western blot was performed using anti-p-Erk antibody. β-actin served as control.

**Supplementary Figure S4 Npac has low enrichment at mouse Pou5f1 genomic region. A**. Schematic diagram at the bottom showed the primer locations at *Pou5f1* genomic region (CR1-4 refer to conserved region 1-4). **B**. Real-time PCR result showed that fold enrichment of Npac ChIP at *Pou5f1* region was lower than 2.

**Supplementary Figure S5 Comparison between Npac genomic distribution with that that of several histone modifications, histone modifiers and ESC-enriched transcription factors. A**. Average whole genome profiles of Npac showed completely opposite to that of H3K4me2. **B**. Representatives heatmap of Npac and histone modifications (H3K9me3, H3K27me3 and H3K4me3). **C**. Representatives heatmap of Npac and histone modifiers of H3K9me3, H3K27me3 and H3K4me3: Eset, EZH2 and MLL2. **D**. Representatives heatmap of Npac and ESC-enriched transcription factors (Nanog, Oct4 and Sox2).

**Supplementary Figure S6 Genomic distribution of Npac and histone H3K36me3 in up-regulated genes upon Npac knockdown showed very low ChIP-seq signal**. Npac and H3K36me3 ChIP-seq peaks at (**A**) developmental genes (*Csf1, Dkk1, Cryab, Hspb2, Wisp1* and *Gata3*), (**B**) MAPK pathway related genes (*Jun, Fas, Egfr* and *Map3k8*), (**C**) Cell death related genes (*Mmd, Cd28, Cstb* and *Krt8*) were shown.

**Supplementary Figure S7 Differences of the binding of RNA Pol II at different gene regions**. The binding of RNA Pol II (**A**) and RNA Pol II Ser2 (**B**) at gene bodies of Utrn were not affected in Npac depleted cells compared to control. The binding of RNA Pol II Ser5 at gene bodies of Nanog (**C**) and Rif1 (**D**) were not significantly affected upon Npac depletion.

**Supplementary Table S1 Sequences of used primers**.

**Supplementary Table S2 Global expression gene changes upon *Npac* RNAi** (cutoff at 1.5 fold change)

**Supplementary Table S3 Gene ontology analysis of altered genes upon *Npac* RNAi** (p<0.05)

**Supplementary Table S4 Global genomic sites of Npac in mESCs**.

**Supplementary Table S5 Gene ontology analysis of Npac ChIP-seq targets**.

