## Supplementary figures and images for "Npac Is a Co-factor of Histone H3K36me3 and Regulates Transcriptional Elongation in Mouse ES Cells"

### Supplentatry figures

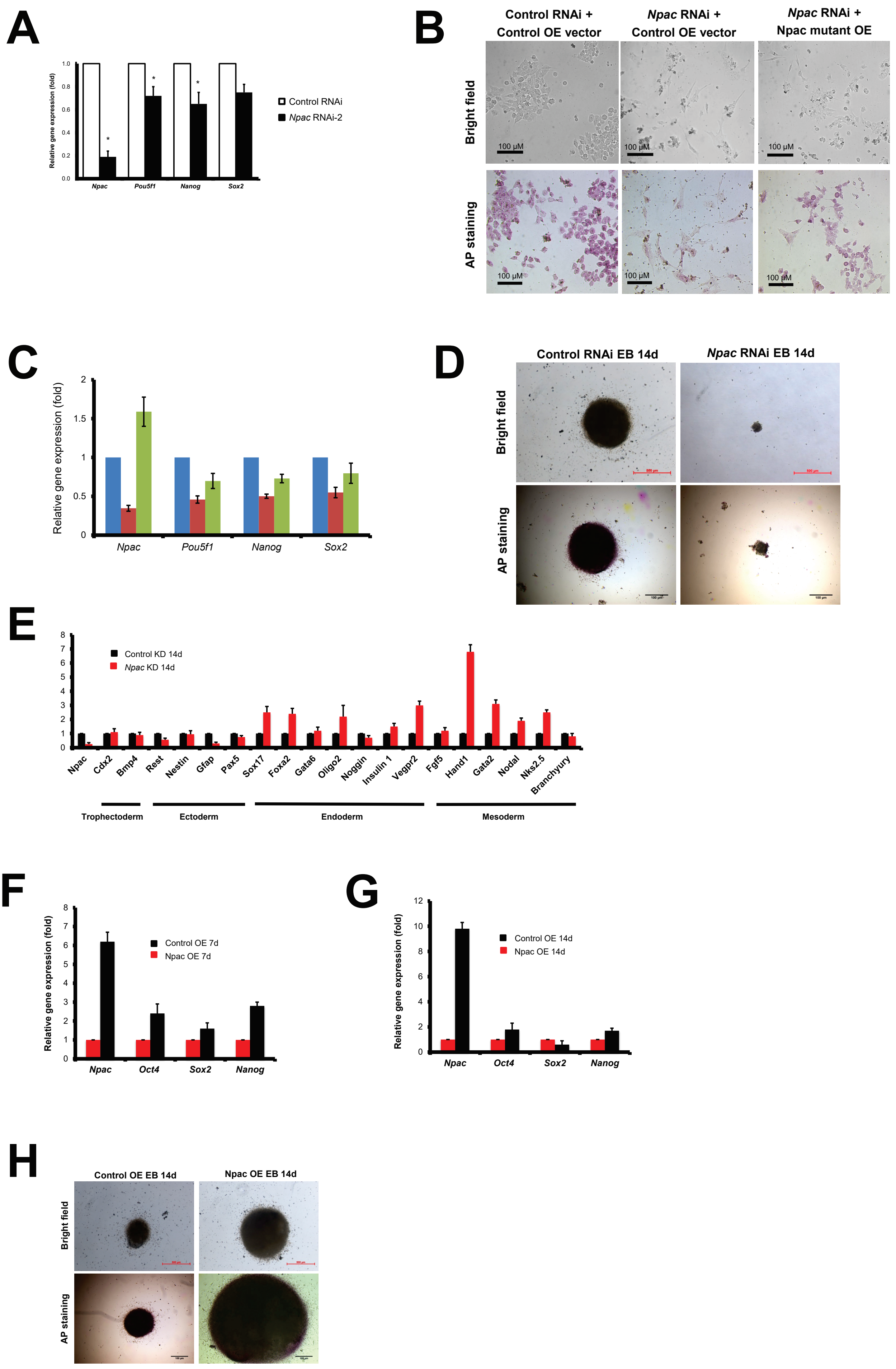

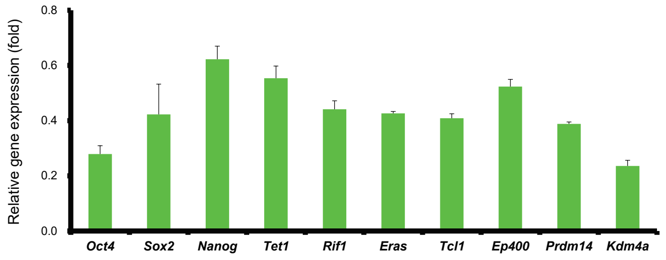

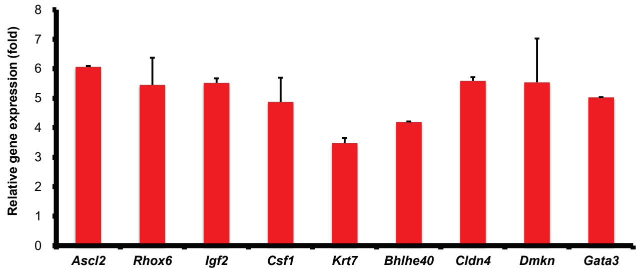

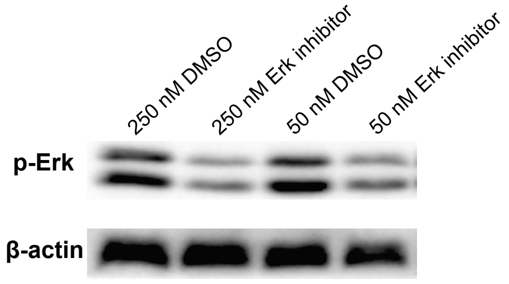

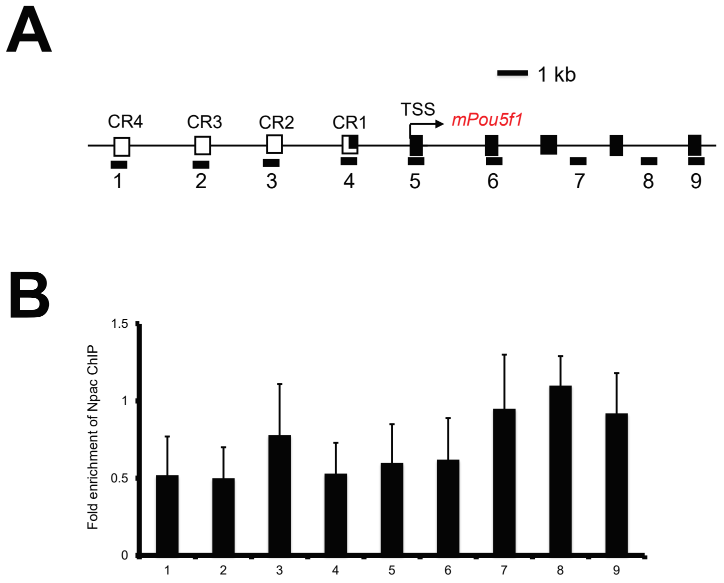

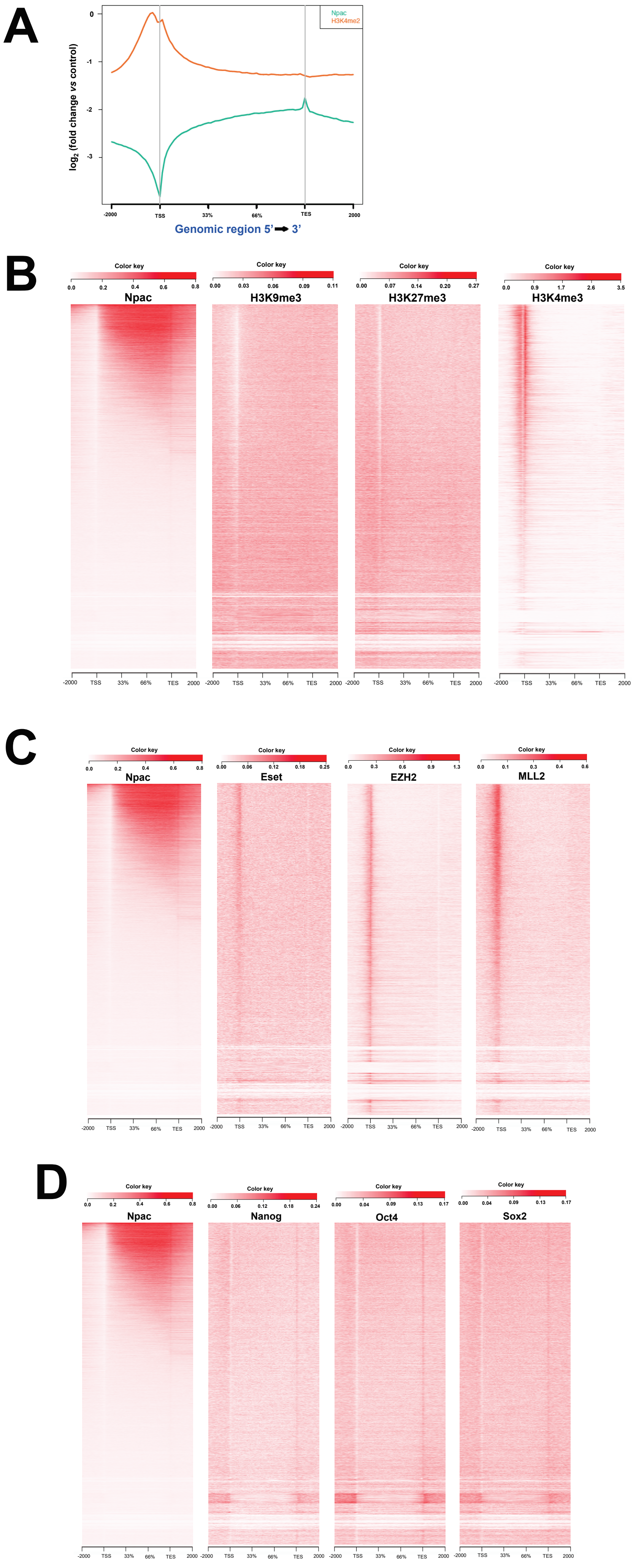

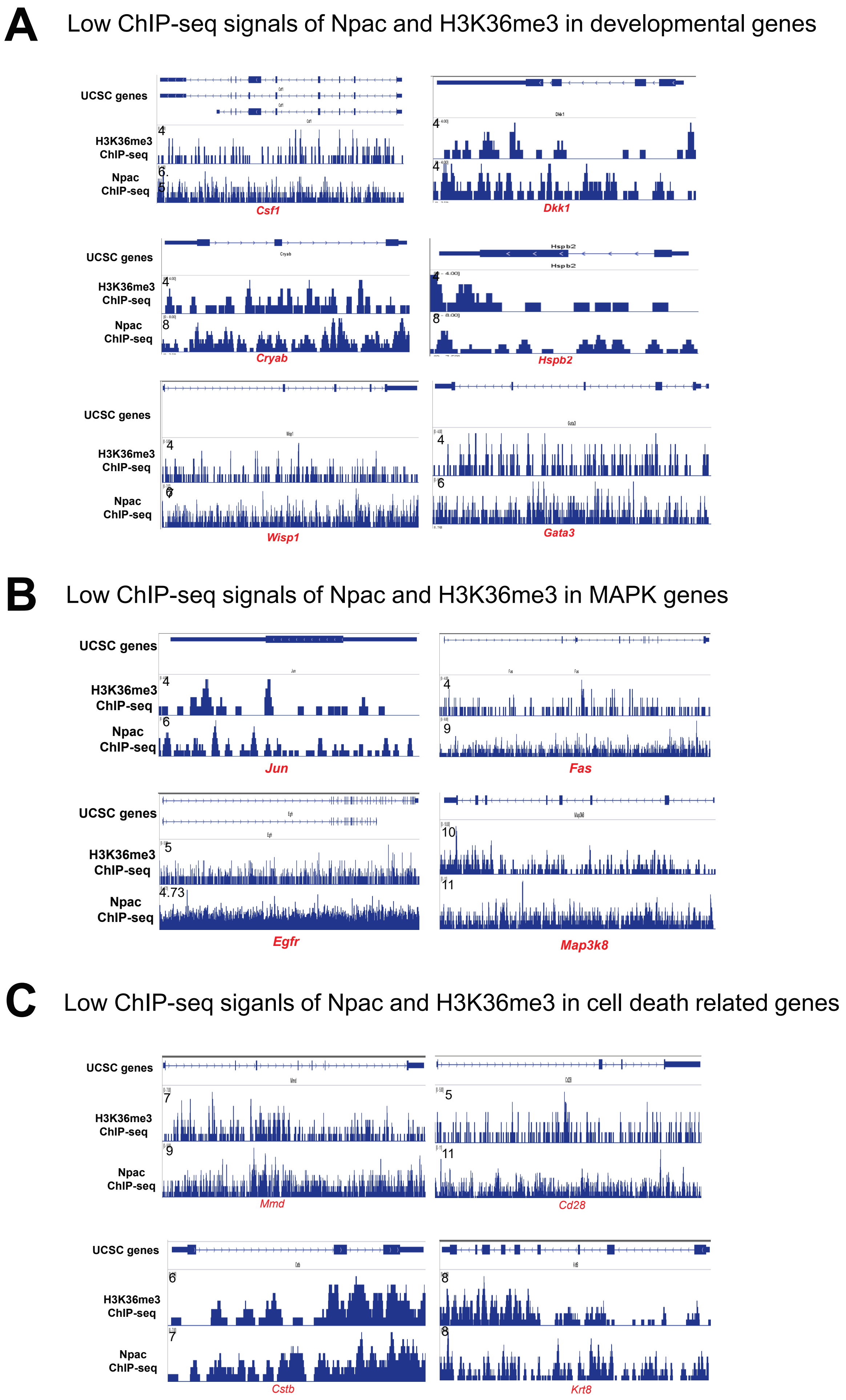

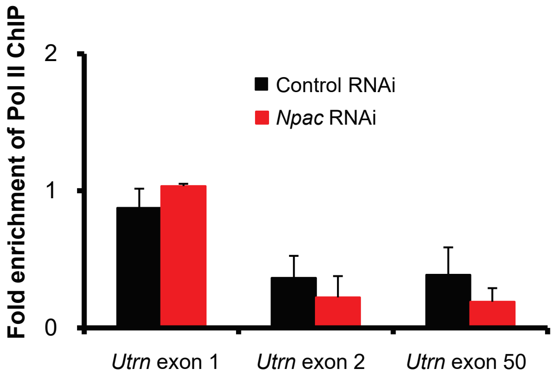

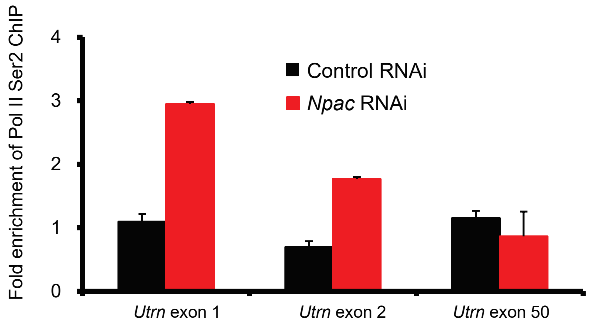

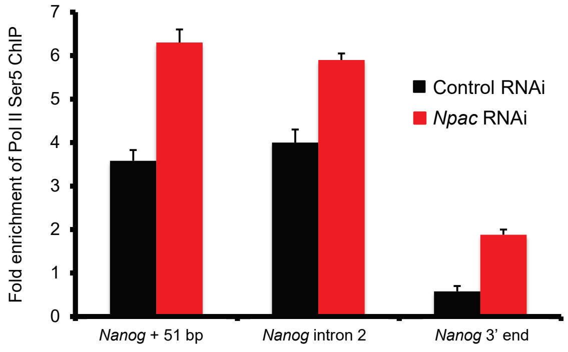

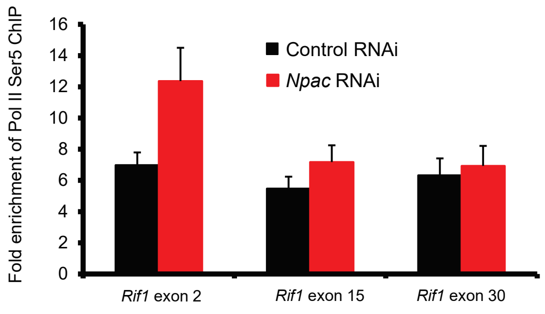
